# Structure of mycobacterial CIII_2_CIV_2_ respiratory supercomplex bound to the tuberculosis drug candidate telacebec (Q203)

**DOI:** 10.1101/2021.07.06.451285

**Authors:** David J. Yanofsky, Justin M. Di Trani, Sylwia Krol, Rana Abdelaziz, Stephanie A. Bueler, Peter Imming, Peter Brzezinski, John L. Rubinstein

## Abstract

The imidazopyridine telacebec, also known as Q203, is one of only a few new classes of compounds in more than fifty years with demonstrated antituberculosis activity in humans. Telacebec inhibits the mycobacterial respiratory supercomplex composed of complexes III and IV (CIII_2_CIV_2_). In mycobacterial electron transport chains, CIII_2_CIV_2_ replaces canonical CIII and CIV, transferring electrons from the intermediate carrier menaquinol to the final acceptor, molecular oxygen, while simultaneously transferring protons across the inner membrane to power ATP synthesis. We show that telacebec inhibits the menaquinol:oxygen oxidoreductase activity of purified *Mycobacterium smegmatis* CIII_2_CIV_2_ at concentrations similar to those needed to inhibit electron transfer in mycobacterial membranes and *Mycobacterium tuberculosis* growth in culture. We then used electron cryomicroscopy (cryoEM) to determine structures of CIII_2_CIV_2_ both in the presence and absence of telacebec. The structures suggest that telacebec prevents menaquinol oxidation by blocking two different menaquinol binding modes to prevent CIII_2_CIV_2_ activity.

## Introduction

Numerous bacteria from the strictly-aerobic genus *Mycobacterium* are human pathogens. In particular, infection by *Mycobacterium tuberculosis* and closely related species result in the disease tuberculosis (TB). In most years TB is the leading cause of death by infectious disease internationally, with an increasing incidence of drug-resistant infections (*Global Tuberculosis Report*, 2020). Nontuberculosis mycobacterial pathogens include *M. leprae*, which causes leprosy, *M. ulcerans*, which causes Buruli ulcer, and *M. avium* and *M. abscessus*, which infect immunocompromised and cystic fibrosis patients, respectively. The discovery of bedaquiline from a phenotypic screen with non-pathogenic *M. smegmatis*, and its subsequent development into an effective therapeutic, has revolutionized the treatment of multidrug-resistant and extensively drug-resistant TB (*Global Tuberculosis Report*, 2020; World Health Organization, 2019). Bedaquiline binds the membrane region of mycobacterial ATP synthase (Andries et al., 2005; Guo et al., 2021; Preiss et al., 2015), blocking proton translocation and ATP synthesis.

Thus, in addition to providing an invaluable therapeutic tool, bedaquiline established oxidative phosphorylation as a target space for antibiotics against mycobacteria. Subsequent to the discovery of bedaquiline, numerous compounds have been identified that target either ATP synthase or the electron transport chain complexes that establish the transmembrane proton motive force (PMF) that drives ATP synthesis (Cook et al., 2017).

In canonical mitochondrial electron transport chains, Complexes I and II oxidize NADH (the reduced form of nicotinamide adenine dinucleotide) and succinate, respectively. The electrons from these substrates are used to reduce a membrane-bound pool of ubiquinone (UQ) to ubiquinol (UQH_2_). Electrons from UQH_2_ are then passed successively to CIII (also known as cytochrome [cyt.] *bc*_1_,), soluble cyt. *c*, and CIV (also known as cyt. *c* oxidase or cyt. *aa*_3_) before ultimately reducing molecular oxygen to water. Complexes I, III, and IV couple electron transfer to proton transfer across the membrane, thereby generating the PMF that drives ATP synthesis. In contrast to mitochondria and many bacteria, mycobacteria possess a branched electron transport chain (Reviewed in (Cook et al., 2017)). Rather than UQ, mycobacterial respiration relies on menaquinone (MQ) as an intermediate electron carrier. MQH_2_ can reduce molecular oxygen via two MQH_2_:O_2_ oxidoreductases: cyt. *bd* and cyt. *bcc-aa*_3_, the latter being equivalent to a combination of canonical CIII and CIV with the stoichiometry CIII_2_CIV_2_. Bioenergetically, the CIII_2_CIV_2_ supercomplex and cyt. *bd* branches of the mycobacterial electron transport chain are not equivalent, with CIII_2_CIV_2_ transferring three protons across the membrane for each electron used to reduce O_2_ while cyt. *bd* transfers only one proton across the membrane for each electron. However, despite this difference, enzyme utilization in mycobacterial electron transport chains can adapt to changes in environmental conditions and treatment with respiratory complex inhibitors (Arora et al., 2014; Berney and Cook, 2010), which complicates targeting of respiration by antimycobacterial drugs (Beites et al., 2019).

Structural analysis of the CIII_2_CIV_2_ supercomplex from *M. smegmatis* led to the discovery that a dimeric type-C superoxide dismutase (SOD) is an integral component of the assembly (Gong et al., 2018; Wiseman et al., 2018). The SOD dimer is found on the periplasmic side of CIII_2_CIV_2_ and is held in place by its two N-terminal tails, which bind to the complex’s QcrB subunits. Both QcrB and the SOD subunit are highly conserved between *M. smegmatis* and *M. tuberculosis*, with 99.6 % and 95 % sequence similarity and 81.1% and 64.8% sequence identity, respectively, suggesting that a similar association occurs in the *M. tuberculosis* enzyme. *M. tuberculosis* mutants that lack SOD are susceptible to killing within macrophages (Piddington et al., 2001), emphasizing the importance of the subunit. Given its position within CIII_2_CIV_2_, it is possible that the SOD subunit abstracts electrons from reactive oxygen species formed during respiration or generated by host-defense mechanisms in the phagolysosome upon phagocytosis of *M. tuberculosis* and directs them through the respiratory chain to contribute to the PMF and ATP synthesis (Gong et al., 2018; Wiseman et al., 2018).

Although killing of *M. tuberculosis* with electron transport chain inhibitors may require simultaneously blocking both the CIII_2_CIV_2_ and cyt. *bd* branches for oxygen reduction (Arora et al., 2014; Beites et al., 2019; Matsoso et al., 2005), high-profile candidate TB therapeutics have been identified that bind to CIII within CIII_2_CIV_2_ (Pethe et al., 2013; Rybniker et al., 2015). Similarly, while CIII_2_CIV_2_ is not essential in *M. smegmatis*, its disruption causes severe growth defects (Matsoso et al., 2005). In contrast, CIII_2_CIV_2_ is essential in *M. leprae* and *M. ulcerans*, which lack the cyt. *bd* oxidase branch of the electron transport chain entirely (Cole et al., 2001; Demangel et al., 2009).

Rather than pumping protons, CIII contributes to the PMF by separating positive and negative charges across the membrane through a mechanism known as the Q cycle (reviewed in (Sarewicz and Osyczka, 2015; Xia et al., 2013). Each CIII contains two sites where redox reactions with MQ occur: a Q_P_ site near the positive (periplasmic) side of the membrane where MQH_2_ is oxidized and a Q_N_ site near the negative (cytoplasmic) side of the membrane where MQ is reduced. In the current understanding of the Q cycle in mycobacteria, oxidation of MQH_2_ in the Q_P_ site leads to release of two protons to the positive side of the membrane. The first electron from this oxidation reaction is passed to a [2Fe–2S] Rieske center (FeS) in subunit QcrA where it consecutively reduces the cyt. *cc* domain of the QcrC subunit and the Cu_A_ di-nuclear center of CIV. The second electron from the MQH_2_ in the Q_P_ site is transferred first to heme *b*_L_ and then heme *b*_H_, both in subunit QcrB of CIII, before reducing a MQ molecule in the Q_N_ site to MQ^•-^. Oxidation of a second MQH_2_ in the Q_P_ site and repetition of this series of events leads to reduction of MQ^•-^ to MQH_2_ in the Q_N_ site, upon abstraction of two protons from the negative side of the membrane, thereby contributing to the PMF and the pool of reduced MQH_2_ in the membrane. Within CIV, electrons are transferred from the Cu_A_ di-nuclear center to the heme *a*_3_-Cu_B_ binuclear catalytic site via heme *a*, driving proton pumping across the membrane.

Telacebec (also known as Q203), was identified in a screen of macrophages infected with *M. tuberculosis* (Pethe et al., 2013). Generation of resistance mutants bearing T313I and T313A mutations in the *qcrB* gene indicated that telacebec targets CIII of the CIII_2_CIV_2_ supercomplex. A recent Phase 2 clinical trial demonstrated a decrease in viable mycobacterial sputum load with increasing dose of telacebec, supporting further development and making telacebec one of only a few new drug classes in more than fifty years with demonstrated antituberculosis activity in humans (van Niekerk et al., 2020). Telacebec may also have clinical utility in treating nontuberculosis mycobacterial infections, such as Buruli ulcer (Almeida et al., 2020; Van Der Werf et al., 2020).

Here we use electron cryomicroscopy (cryoEM) to investigate how telacebec inhibits purified CIII_2_CIV_2_ from *M. smegmatis*. We develop conditions for CIII_2_CIV_2_ activity assays that limit the spontaneous autoxidation of MQH_2_ analogues that has hampered previous analysis of MQH_2_:O_2_ oxidoreductase activity with purified CIII_2_CIV_2_. The assays show that telacebec inhibits CIII_2_CIV_2_ activity at concentrations comparable to those that inhibit electron transfer in mycobacterial membranes. CryoEM of CIII_2_CIV_2_ demonstrates both the presence of the LpqE subunit (Gong et al., 2018) and different conformations of the cyt. *cc* domain (Wiseman et al., 2018), which have previously been observed separately but not together. 3D variability analysis of the structure shows that the SOD subunit can move toward cyt. *cc*, supporting the possibility of direct electron transfer from superoxide to CIV. CryoEM of the CIII_2_CIV_2_:telacebec complex allows localization of the telacebec binding site with the imidazopyridine moiety and A-benzene ring of telacebec forming most protein-inhibitor contacts.

## Results

### Structure of CIII_2_CIV_2_ reveals movement of SOD subunit and cyt. cc domain

In order facilitate isolation of CIII_2_CIV_2_, we used the ORBIT (oligonucleotide-mediated recombineering followed by Bxb1 integrase) strategy (Murphy et al., 2018) to introduce sequence for a 3’FLAG affinity tag into the chromosomal DNA of *M. smegmatis* immediately 3¢ to the *qcrB* gene. While *M. smegmatis* is typically grown in 7H9 medium supplemented with albumin, dextrose, and sodium chloride (ADS), we found that supplementing instead with tryptone, dextrose, and sodium chloride (TDS), which is more economical for large scale culture, gave equivalent or superior growth. Purification of CIII_2_CIV_2_ from *M. smegmatis* grown in these conditions gave a high yield of enzyme with clear bands for most of the known subunits of the complex (**Fig. 1A**). We observed that following affinity purification, gel filtration chromatography of the enzyme led to depletion of the LpqE and SOD subunits (**Fig. 1A**, ***right***) compared to affinity purification alone (**Fig. 1A**, ***left***), and consequently this purification step was avoided.

**Figure 1.**
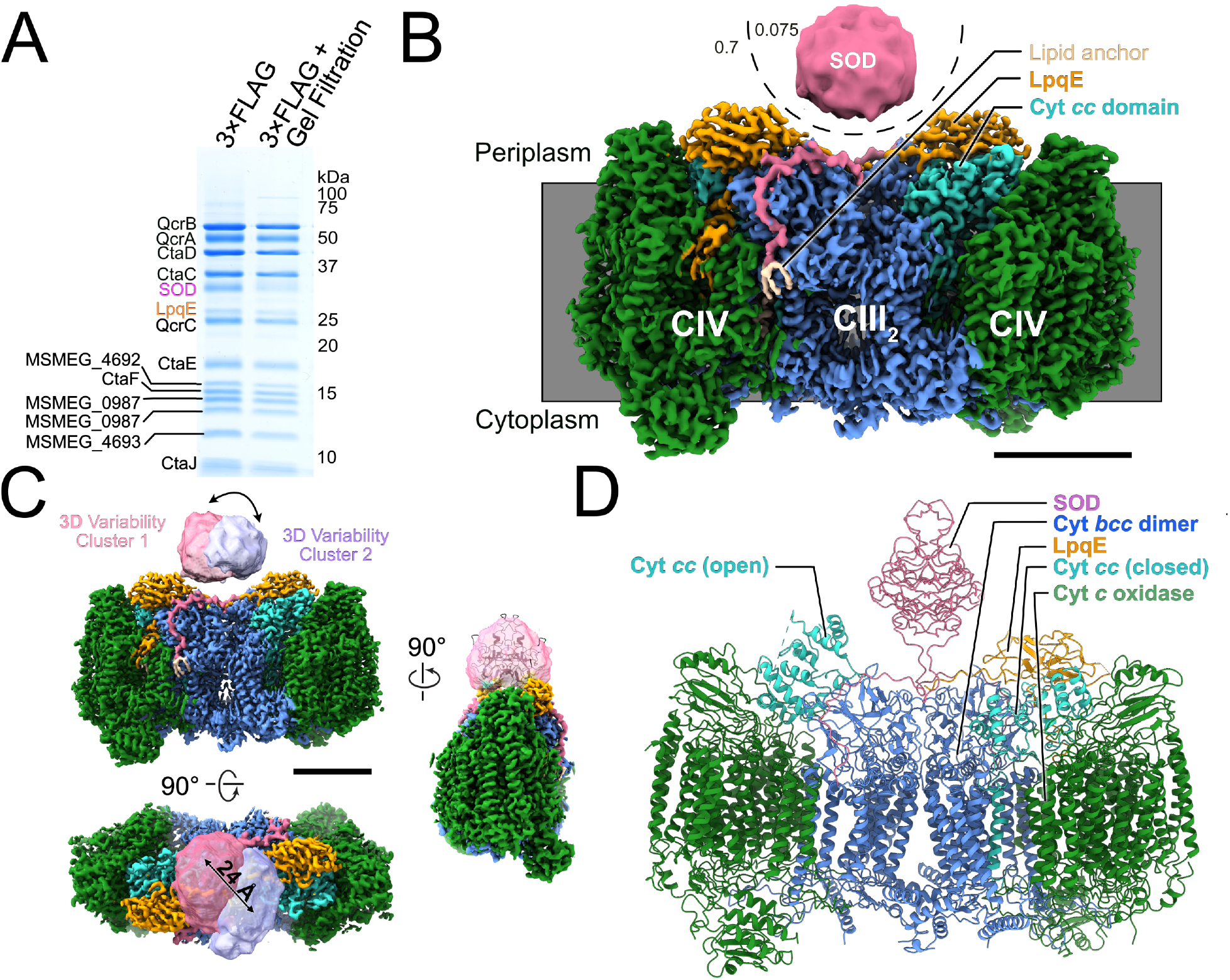
Structure of the *M. smegmatis* CIII_2_CIV_2_ respiratory complex. **A,** SDS-PAGE shows most of the known subunits of the complex and indicates that the SOD and LpqE subunits are depleted by gel filtration chromatography. **B,** CryoEM map of the CIII_2_CIV_2_. The different density thresholds for the SOD subunit and the rest of the complex are indicated. Scale bar, 50 Å. **C,** Three-dimensional variability analysis indicates two different clusters of particle images (“cluster 1” and “cluster 2”) that show the SOD subunit in different positions over the 2-fold symmetry axis of the complex. Scale bar, 50 Å. **D,** An atomic model for the CIII_2_CIV_2_ complex with SOD fitted into the map and showing one half of the complex missing the LpqE subunit and with the cyt. *cc* domain in the ‘open’ conformation and the other half of the complex possessing the LpqE subunit and with the cyt. *cc* domain in the ‘closed’ conformation.

CryoEM of the CIII_2_CIV_2_ preparation allowed for calculation of a 3D map of the enzyme at a nominal resolution of 3.0 Å (**Fig. 1B**, **Figure 1 – Figure Supplement 1**, and **Table 1**). The map shows strong density for the LpqE subunit (**Fig. 1B**, ***orange***). LpqE was observed in one previous structural study of CIII_2_CIV_2_ from *M. smegmatis* (Gong et al., 2018) but was absent in another (Wiseman et al., 2018), presumably due to depletion of the subunit during purification of the supercomplex. In the structure missing LpqE, the cyt. *cc* domain of subunit QcrC adopts both an ‘open’ and a ‘closed’ conformation, while the structure with LpqE was found only in the closed conformation. The closed conformation creates a direct electronic connection between heme *c*_II_ of CIII and Cu_A_ of CIV (Gong et al., 2018; Wiseman et al., 2018). In the open conformation, heme *c*_II_ from the cyt. *cc* domain is too far from Cu_A_ to allow electron transfer, leading to the hypothesis that switching between the closed and open conformations plays a role in controlling the flow of electrons through the supercomplex (Wiseman et al., 2018). In contrast, LpqE was hypothesized to strengthen the physical attachment between CIII and CIV (Gong et al., 2018). 3D variability analysis with the current dataset (Punjani and Fleet, 2021), focused on one half of the supercomplex, revealed complexes with and without LpqE. Where LpqE was missing, the cyt. *cc* domain exhibits the open conformation, while complexes with LpqE show only the closed conformation of cyt. *cc* (**Figure 1 – Figure Supplement 2**). Clashes between LpqE and the open conformation of cyt. *cc* suggest that LpqE prevents the open conformation.

**Table 1.** CryoEM structure determination.

**A. Cryo-EM data acquisition and image processing.**
| **Data Collection** |  |
| --- | --- |
| Electron Microscope | Titan Krios G3 |
| Camera | Falcon 4 |
| Voltage (kV) | 300 |
| Nominal Magnification | 75,000 |
| Calibrated pixel size (Å) | 1.03 |
| Total exposure (e/Å<sup>2</sup>) | 43.5 |
| Exposure rate (e/pixel/s) | 5.99 |
| Number of exposure fractions | 29 |
| Defocus range (µm) | 0.7–2 |
| **Image Processing** |  |
| Motion correction software | cryoSPARC v3 |
| CTFestimation software | cryoSPARC v3 |
| Particle selection software | cryoSPARC v3 |
| Micrographs used in inhibitor-free dataset | 4,308 |
| Micrographs used in telacebec-bound dataset | 2,793 |
| Particle images selected in inhibitor free dataset | 1,037,709 |
| Particle images selected in telacebec-bound dataset | 387,777 |
| 3D map classification and refinement software | cryoSPARC v3 |

**B. Model statistics.**
| **Dataset** | **Inhibitor-free** | **Telacebec-bound** |
| --- | --- | --- |
| **Associated PDB ID** |  |  |
| **Modelling and refinement software** | Coot, phenix, ISOLDE | Coot, phenix, ISOLDE |
| **Protein residues** | 6,058 | 6,075 |
| **Ligand** | 9XX: 4, 9Y0: 6, 9YF: 8, FES: 2, HEC: 4, HEA: 4, MQ9: 10, HEM: 4, PLM: 4, CU: 8 | 9XX: 4, 9Y0: 6, 9YF: 8, FES: 2, HEC: 4, HEA: 4, MQ9: 10, HEM: 4, PLM: 4, CU: 8, QTE: 1 |
| **RMSD bond length (Å)** | 0.005 | 0.006 |
| **RMSD bond angle (°)** | 1.485 | 1.63 |
| **Ramachandran outliers (%)** | 0.22 | 0.52 |
| **Ramachandran favoured (%)** | 92.58 | 92.67 |
| **Rotamer outliers (%)** | 0 | 0 |
| **Clash score** | 19.29 | 25.01 |
| **MolProbability score** | 2.25 | 2.35 |
| **EMRinger score** | 3.12 | 2.58 |

In previous studies the SOD subunit of CIII_2_CIV_2_ was poorly resolved in cryoEM maps and appeared as a diffuse density (Gong et al., 2018; Wiseman et al., 2018). In the present map the overall shape of SOD, although still at lower density than the rest of the complex, was more readily apparent (**Fig. 1B**, ***pink***). The N-terminal anchors from the SOD dimer that bind to subunit QcrB are well resolved and terminate at the middle of the complex with a lipid anchor (**Fig. 1B**, ***beige***). The improved density for the SOD subunit allowed fitting of a homology model of the protein into the map with a slight rotation relative to how it was fit previously (**Fig. 1C**, ***right***). 3D variability analysis (**Fig. 1C**) shows that SOD moves between the center of the complex, where it was observed previously (Gong et al., 2018; Wiseman et al., 2018) to immediately above heme *c*I. This proximity suggests that SOD may indeed transfer electrons abstracted from superoxide in the periplasm of *M. smegmatis* to CIV thereby contributing to the PMF (Gong et al., 2018; Wiseman et al., 2018), although this hypothesis requires further testing. The overall resolution of the map, which is somewhat better than in previous studies, allowed refinement of an atomic model for CIII_2_CIV_2_ including residues T82, E83, A123 to D131, and S183 from LpqE, and residues H57 to G78 from MSMEG_4693 (also known as CtaJ), which could not be modelled previously (**Fig. 1D**, **Fig. 1 – Figure Supplement 3**, and **Table 1**). The model shown in **Fig. 1D** illustrates one cyt. *cc* domain in the closed position with LpqE present (**Fig. 1D**, ***right side: cyan and orange***) and the other cyt. *cc* domain in the open position without LpqE (**Fig. 1D**, ***left side: cyan***).

### Nanomolar telacebec inhibits oxidoreductase activity with purified CIII_2_CIV_2_

To investigate inhibition of CIII_2_CIV_2_ by telacebec, we established a supercomplex activity assay, based on measurement of oxygen consumption with a Clark-type electrode. The mycobacterial electron transport chain uses MQH_2_ as the electron donor for CIII_2_CIV_2_ while in canonical mitochondrial electron transport chains UQH_2_ donates electrons to CIII_2_ (Cook et al., 2017). Both UQH_2_ and MQH_2_ are insoluble in aqueous solution and consequently soluble analogues must be employed as substrates in assays with detergent-solubilized enzymes. The midpoint potentials of the redox sites of mycobacterial CIII_2_ are lower than those of canonical mitochondrial CIII_2_ (Kao et al., 2016), and as a result UQH_2_ analogues typically used in CIII_2_ assays are not able to reduce CIII_2_ of the *M. smegmatis* supercomplex. MQH_2_ analogues capable of reducing CIII_2_CIV_2_ suffer from autoxidation at neutral or basic pH, which leads to oxygen reduction even in the absence of enzyme. This background oxygen-reduction rate is typically subtracted from that observed in the presence of enzyme to calculate the enzyme-catalyzed oxidoreductase activity. Previous measurement of inhibition of CIII_2_CIV_2_ by telacebec employed menadiol as the electron donor, which is highly soluble in aqueous solution, but rapidly autoxidizes, leading to the background signal from autoxidation in the absence of enzyme being higher than the signal from oxygen reduction by the enzyme (Gong et al., 2018). Instead, we selected 2,3-dimethyl-[1,4]naphthoquinone (DMW), which we chemically reduced to DMWH_2_, as the substrate for oxygen reduction assays (Graf et al., 2016; Wiseman et al., 2018). Initial activity assays led to anomalous results where addition of low concentrations of CIII_2_CIV_2_ to the assay mixture appeared to decrease the rate of oxygen reduction below the background autoxidation rate. On subsequent investigation, we realized that at low concentrations of CIII_2_CIV_2_ the SOD subunit suppresses autoxidation of DMW more than CIII_2_CIV_2_ catalyzes oxidation of DMW. This suppression of QH2 autoxidation by SOD, which has been described previously (Cadenas et al., 1988), can lead to apparent negative activities for CIII_2_CIV_2_ when the background autoxidation is subtracted. To remove this source of error, we established that bovine C-type SOD can similarly limit the autoxidation of DMW (**Fig. 2A**, ***blue* and *orange* curves**). Thus, by adding an excess of exogenous bovine SOD to assays, the CIII_2_CIV_2_’s DMW:O_2_ oxidoreductase activity can be measured with suppression of DMW autoxidation (e.g. **Fig. 2A**, ***green***). With 500 nM SOD added, the CIII_2_CIV_2_’s DMW:O_2_ oxidoreductase activity was measured at 91 ±4 e^−^/s (±s.d., n=6 independent assays), which is nearly an order of magnitude greater than the apparent activity found previously (Wiseman et al., 2018). We subsequently added 500 nM bovine SOD to all assays to limit autoxidation of DMW.

**Figure 2.**
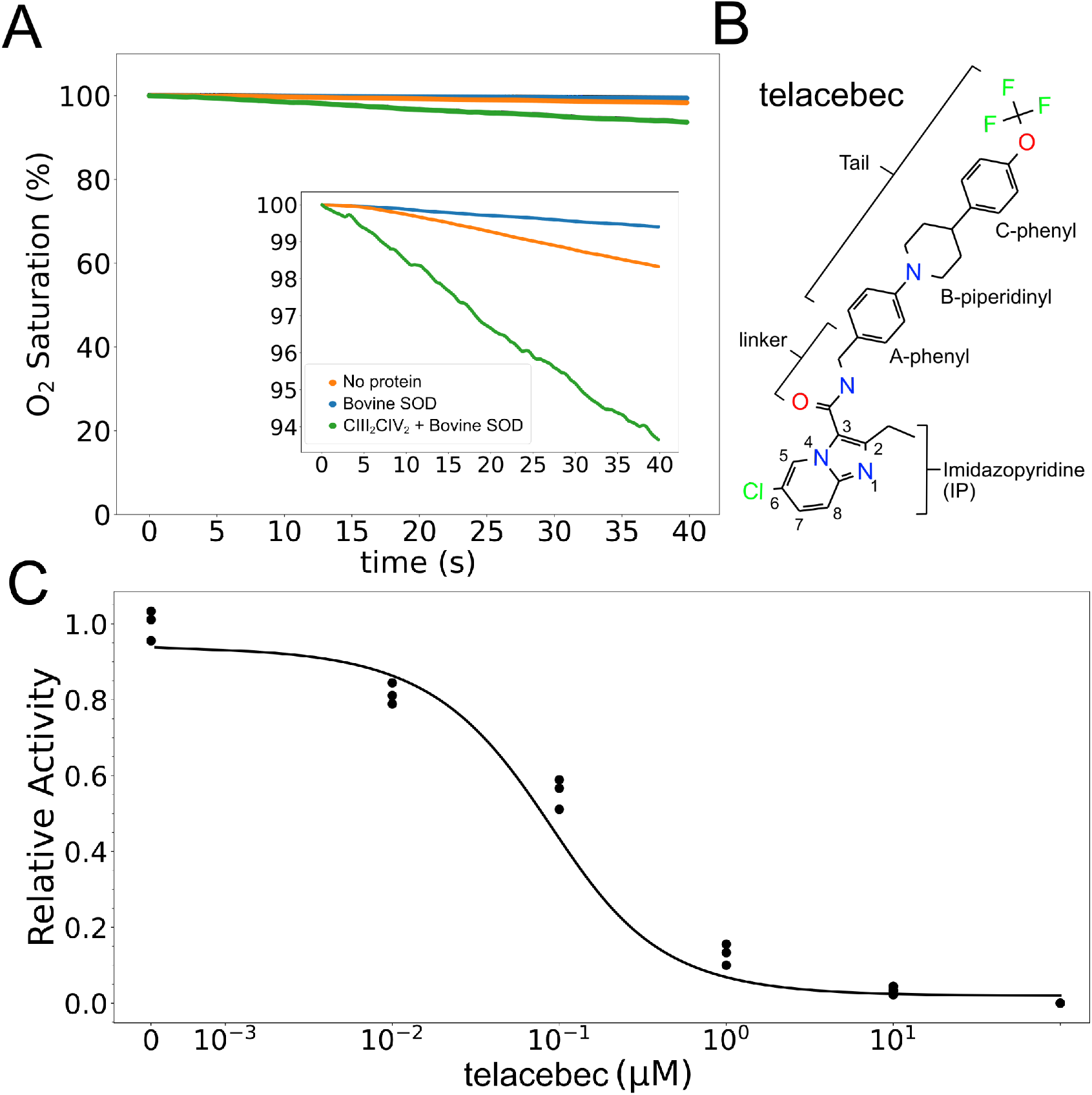
Assay of DMWH_2_:O_2_ oxidoreductase activity by CIII_2_CIV_2_. **A,** An oxygen reduction assay shows that autoxidation of 2,3-dimethyl-[1,4]naphthoquinone (DMW), *blue curve*, is decreased by the presence of 500 nM bovine SOD, *orange curve*. Measurement of oxygen reduction by CIII_2_CIV_2_ in the presence of bovine SOD, *green curve*, allows calculation of CIII_2_CIV_2_ activity. **B,** Structure of CIII_2_CIV_2_ inhibitor telacebec (Q203). **C,** Titration of CIII_2_CIV_2_ (60 nM) with telacebec shows an IC_50_ of 53±19nM (±s.d., n=3 independent titrations) with 100 μM DMWH_2_.

Telacebec (**Fig. 2B**) is a potent inhibitor of mycobacterial CIII_2_ (Pethe et al., 2013). The compound consists of an imidazo[1,2-a]pyridine attached via an amide linker to a N-[(4-{4-[4-(trifluoromethoxy)phenyl]piperidin-1-yl}phenyl)methyl] ‘tail’. Titrations of CIII_2_CIV_2_ activity with varying concentrations of telacebec (**Fig. 2C**) show an IC_50_ of 53 nM ±19(±s.d., n=3 independent titrations) with 65 nM CIII_2_CIV_2_ and 100 μM DMW. This IC_50_ is lower than the 840 ±22nM seen with the menadiol based assay (Gong et al., 2018), but higher than the 20 nM concentration needed to inhibit 50% of respiratory chain activity with inverted membrane vesicles from *M. smegmatis* (Lu et al., 2018) or 2.7 nM required to inhibit the 50% of *M. tuberculosis* growth in liquid culture (Pethe et al., 2013).

The *M. tuberculosis* telacebec resistance mutations T313A and T313I (Pethe et al., 2013), equivalent to mutation of Thr308 in *M. smegmatis*, are near the Q_P_ site and suggest that the inhibitor could interfere with MQ binding to CIII_2_CIV_2_. The increased IC_50_ in the current assay compared to assays with inverted membrane vesicles or bacterial growth in liquid culture may be due to the binding affinity or high concentration of DMW, which could allow DMW to compete with telacebec for binding to the complex.

### The CIII_2_CIV_2_ structure has endogenous MQ in its Q_P_ site

As telacebec is expected to bind near the Q_P_ site of CIII_2_CIV_2_, we carefully characterized this site in the cryoEM map of the enzyme in the absence of inhibitor. The Q_P_ site is near the periplasmic side of the membrane, located between heme *b*_L_ and the FeS cluster (**Fig. 3A**), and is formed by several loops and ahelices from both the QcrB and QcrA subunits (**Fig. 3B**). The arrangement of structural elements in the site is conserved from other CIIIs (Sarewicz and Osyczka, 2015). The entrance to the Q_P_ site is formed by the C and F transmembrane ahelices, and the cd2 ahelix that separates the periplasmic side of the Q_P_ pocket from the QcrA subunit. The ef helix and ef loop from the QcrB subunit are deeper in the Q_P_ site, as is a short section from the QcrA subunit that includes the FeS-bound His368 residue (**Fig. 3B**).

**Figure 3.**
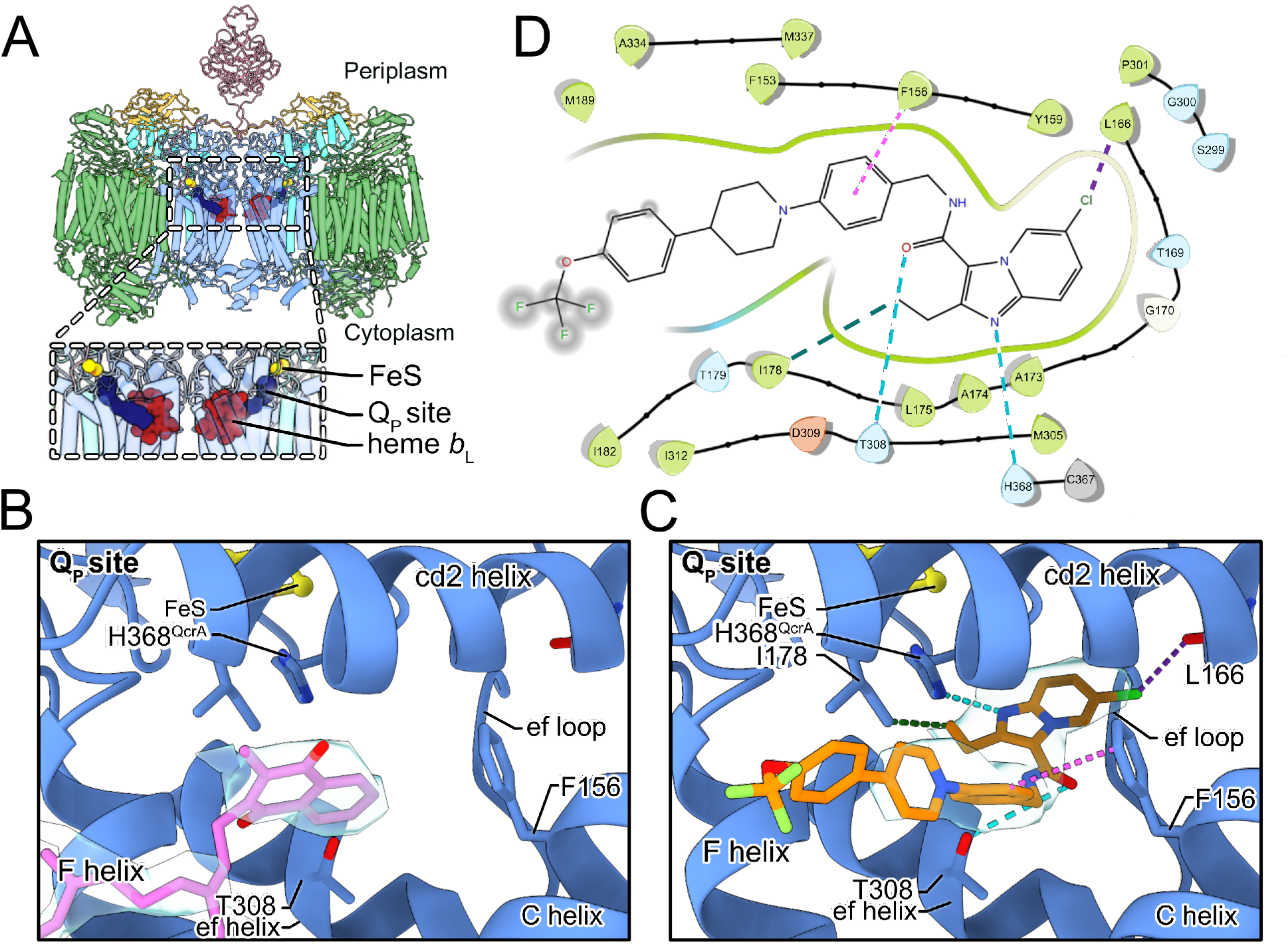
Telacebec binding to the Q_P_ site. **A,** The dashed boxes indicate the two Q_P_ sites in CIII_2_CIV_2_, each showing menaquinone (*blue*), the Rieske protein FeS (*yellow*), and heme *b_L_* (*red*). **B,** In the inhibitor-free structure, there is density for endogenous menaquinone (*pink model* and *grey surface*) distal from the FeS group amongst the well-conserved structural elements of the Q_P_ site. **C,** In the inhibitor-bound structure there is density for telacebec (*orange model* and *grey surface*) deeper in the Q_P_ site where it can form numerous interactions with the protein, including possible hydrogen bonds (*dashed teal lines*), hydrophobic interactions (*dashed green line*), a halogen bond (*dashed purple line*), and an aromatic interaction (*dashed pink line*). **D,** A 2D representation of the interactions between telacebec and residues of CIII_2_CIV_2_ using the same colour convention as in part C.

In the inhibitor-free structure there is strong density for endogenous MQ in the Q_P_ site (**Fig. 3B**, ***pale blue surface***). The naphthoquinone head group of MQ is positioned near the entrance to the site, between the F and C helices, with the MQ tail trailing towards the Q_N_ site (**Fig. 3A**). This position for endogenous MQ was reported in a previous study of CIII_2_CIV_2_ from *M. smegmatis* (Gong et al., 2018). In this position, the naphthoquinone head group is ~14 Å away from the FeS cluster and the hydroxyl proton is ~15 Å from His368, which is too far for rapid coupled electron and proton transfer from MQH_2_ to FeS and His368, respectively. This distance contrasts the deeper binding position adopted by UQ in ovine CIII_2_ (Letts et al., 2019). It is also further from the FeS than the position observed for UQ-analogue inhibitors such as stigmatellin bound to chicken CIII_2_ (Zhang et al., 1998), as well as 5-undecyl-6-hydroxy-4,7-dioxobenzothiazole (Esser et al., 2004) and 2-n-nonyl-4-hydroxyquinoline N-oxide (Gao et al., 2003) bound to bovine CIII_2_. In the deeper position the head groups of UQ or its analogues are wedged between the ef helix/loop, and the cd2 helix, with the tail trailing between the F and C helix at the Q_P_ site entrance (**Fig. 3 – Figure Supplement 1A**). In these structures, the distance between the quinone head group and the FeS cluster depends on the position of the mobile Rieske head domain but can be as little as ~7 Å, allowing for rapid electron transfer from the UQH_2_ to the FeS (Moser et al., 2006).

### Telacebec replaces MQ in the Q_P_ site of active CIII_2_CIV_2_

Our initial attempts to image CIII_2_CIV_2_ with telacebec failed to resolve the inhibitor (**Fig. 3 – Figure Supplement 1B**, ***left***), leading us to consider the possibility that inhibitor binding occurs during substrate turnover by the enzyme. CryoEM of CIII_2_CIV_2_ in the presence of DMWH_2_ but without telacebec confirmed that under these conditions the density in the Q_P_ site was indistinguishable from MQ seen with the enzyme at rest (**Fig. 3 – Figure Supplement 1B**, ***right***). We then incubated CIII_2_CIV_2_ with both DMWH_2_ and telacebec to produce an inhibited complex and determined the structure of this complex to a nominal resolution of 3.0 Å by cryoEM (**Fig. 1 – Figure Supplement 1**, **Table S1**). Telacebec binding did not cause large-scale conformational changes in CIII_2_CIV_2_ but produced a clear density for the inhibitor in each of the two Q_P_ sites in the CIII_2_ dimer (**Fig. 3C**). The inhibitor’s imidazopyridine moiety, amide linker region, and A-phenyl and B-piperidinyl moieties are all resolved clearly, with weaker density toward the end of the tail, which points into the lipid bilayer toward the cytoplasmic side of the membrane (**Fig. 3C**, ***pale blue surface***). These densities show that telacebec binds with its head group deep within the Q_P_ binding pocket in a pose similar to UQ and the UQ-analogue inhibitors bound within the canonical CIII_2_ as described above (**Fig. 3 – Figure Supplement 1A**). Telacebec’s imidazopyridine moiety forms multiple interactions with the protein to stabilize inhibitor binding. Although hydrogen bonds cannot be detected with complete confidence at the present resolution, the position of the N1 nitrogen in telacebec’s imidazopyridine moiety is consistent with formation of a hydrogen bond with the His368 from the QcrA subunit, which also binds the FeS group (**Fig. 3C** **and** **D**, ***dashed teal line***). The occurrence of a similar hydrogen bond between UQ and the equivalent histidine residue in canonical CIII_2_ (Zhang et al., 1998) has been proposed to coordinate the Q cycle (e.g. see (Sarewicz and Osyczka, 2015)). The 2-ethyl group from the imidazopyridine is ~4 Å away from Ile178 from the QcrB subunit, providing hydrophobic interactions (**Fig. 3C** **and** **D**, ***dashed green line***), while the 6-chloro group from the imidazopyridine is close to the backbone carboxyl group of Leu166 from the QcrB subunit, enabling formation of a possible halogen bond (**Fig. 3C** **and** **D**, ***dashed purple line***).

In addition to its imidazopyridine moiety, the amide linker and tail of telacebec also contact subunit QcrB, stabilizing binding. Thr308, which is homologous with *M. tuberculosis* Thr313 and is known to be important for binding (Pethe et al., 2013), is <4 Å away from the linker region of telacebec. Although the rotameric state of Thr308 is ambiguous at the current resolution, one of the rotamer states could form a stabilizing hydrogen bond with the carbonyl group of the linker region (**Fig. 3C** **and** **D**, ***dashed teal line***). Finally, Phe156 is ~3.5 Å from the A-phenyl group of telacebec, allowing for aromatic-aromatic interaction between the protein and inhibitor (Burley and Petsko, 1985) (**Fig. 3C** **and** **D**, ***dashed pink line***) Interestingly, in the inhibitor-free specimen, the MQ head group is positioned similarly to the A-phenyl ring of telacebec and may form similar stabilizing contacts with the QcrB subunit (**Fig. 3B** **and** **C**).

### Telacebec blocks both the standby and oxidation positions for MQ

The telacebec-bound structure also provides insight into the basic mechanism of MQ:O_2_ oxidoreductase activity by CIII_2_CIV_2_. Multiple MQ binding sites were modelled in an earlier 3.5 Å resolution cryoEM density map of CIII_2_CIV_2_, but the significance of these sites was not clear (Gong et al., 2018). The structure presented here shows that telacebec, which can inhibit MQ:O_2_ oxidoreductase activity completely (**Fig. 2C**), binds only at the Q_P_ site. Therefore, it is unlikely that MQ-binding other than within the Q_P_ site is involved in electron transfer to oxygen. During the Q cycle, two molecules of MQH_2_ are oxidized to MQ at the Q_P_ site for each molecule of MQ reduced to MQH_2_ at the Q_N_ site. It is possible that MQ is channeled between the Q_P_ and Q_N_ sites by staying loosely bound to the supercomplex surface, with the additional MQ sites serving as intermediate positions along the channeling pathway. A similar model was suggested for the spinach *b*_6_*f* complex, which is structurally and functionally related to CIII and carries out the Q cycle in plant chloroplasts with the hydrophobic electron carrier plastiquinone (Malone et al., 2019). It is also possible that these alternative MQ sites sequester MQ, increasing the local concentration of substrate near its two binding sites, which has been seen to increase the local concentration of ligands in other systems (Vauquelin and Charlton, 2010).

Within the Q_P_ site different positions for both UQ and MQ have been described previously, both for canonical CIII and for CIII within a CIII_2_CIV_2_ supercomplex, respectively (Ding et al., 1992; Moe et al., 2021). As discussed above, in the present structure, the endogenous MQ is too far for rapid transfer of protons and electrons from MQH_2_ to His368 and the FeS center, respectively (**Fig. 4A**). This binding site within Q_P_ is known as the Q1b position. In contrast, movement of MQ deeper within the Q_P_ site to the Q1a position would bring it less than 10 Å from FeS, close enough to donate electrons to the redox center and protons to His368 (**Fig. 4B**). In an emerging model for CIII_2_CIV_2_ function, the Q1b position serves as a “stand-by” site for MQ, with oxidation of the substrate occurring only upon relocation to Q1a (see also (Moe et al., 2021; Mulkidjanian, 2005)). The structure of CIII_2_CIV_2_ with telacebec bound shows how that the compound serves as dual site inhibitor (**Fig. 4C**), with the imidazopyridine group bound to the Q1a position and A-phenyl portion of the tail bound to the Q1b position. Indeed, the possible hydrogen bond between Thr308 and the carboxyl group in the linker region of telacebec could also occur between Thr308 and MQ, stabilizing it in the Q1b position. Therefore, the observed pose of the inhibitor within the enzyme not only blocks MQ access to the FeS center but fills the Q_P_ site entirely.

**Figure 4.**
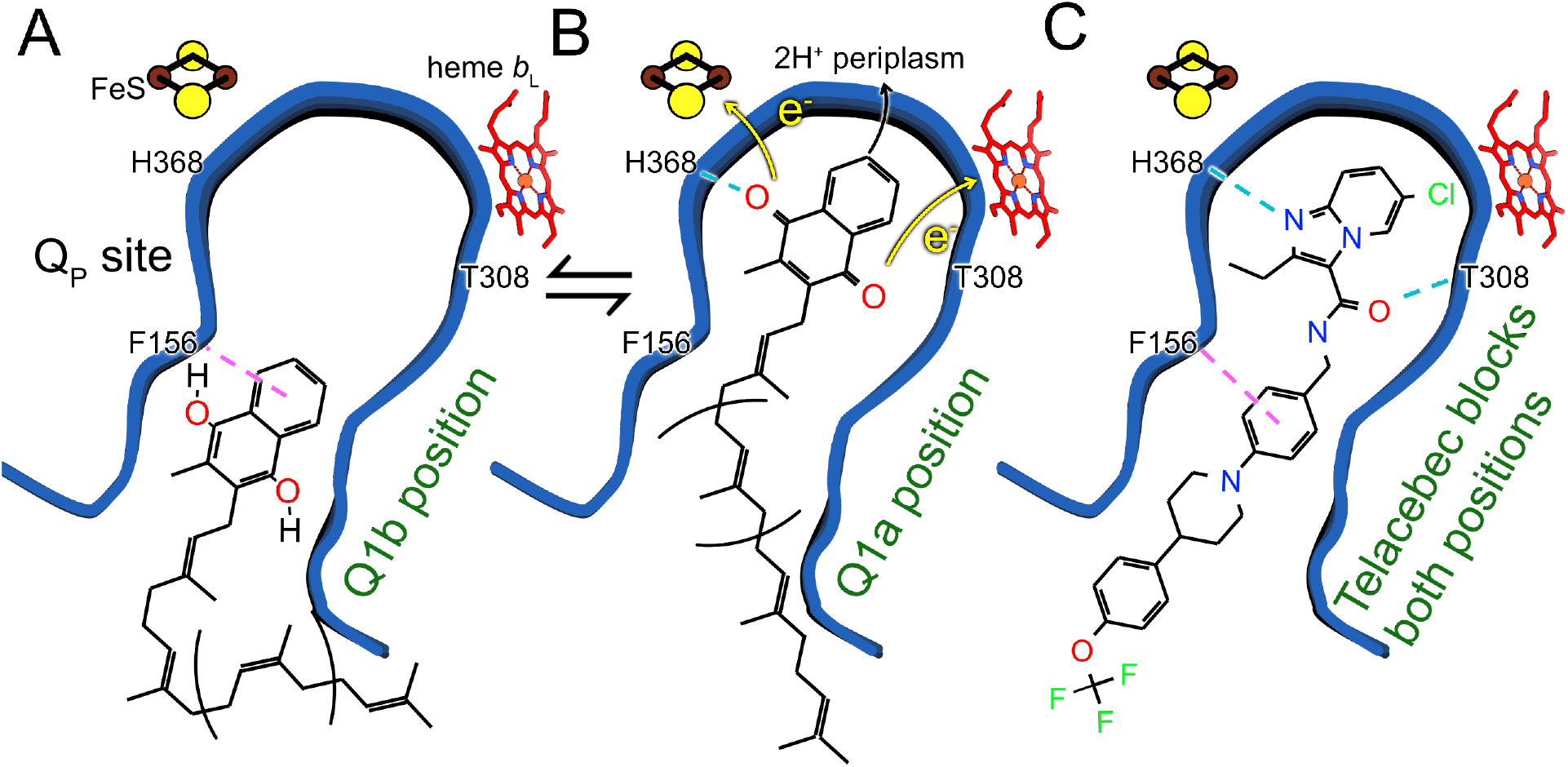
Model for oxidation of MQH_2_ in the Q_P_ site and how telacebec blocks it. An emerging model for MQH_2_ reduction at the Q_P_ site proposes that the substrate binds in the Q1b position where it is too far from FeS to donate protons and electrons (**A**). Upon moving deeper into the Q_P_ site to the Q1a position, MQH_2_ is oxidized to MQ, donating its first electron to FeS, its second electron to heme *b*_L_, and releasing two protons to the positively charged periplasmic side of the lipid bilayer (**B**). Telacebec binds deep within the Q_P_ site, forming numerous interactions with the protein and blocking both the Q1a and Q1b positions (**C**).

Inhibiting CIII_2_CIV_2_ has demonstrated antituberculosis activity in humans (van Niekerk et al., 2020), even though *M. tuberculosis* possesses cyt. *bd* as an alternative enzyme that can oxidize MQH_2_ and sustain the electron flux to oxygen in the mycobacterial membrane. More potent killing of mycobacterial pathogens can be accomplished by simultaneous inhibition of both the CIII_2_CIV_2_ and cyt. *bd* terminal oxidases (Beites et al., 2019). The present study demonstrates how cryoEM can reveal the mechanisms of electron transport chain inhibitors, enabling new strategies for targeting mycobacterial infections.

## Methods

### Construction of M. smegmatis strain, cell culture, and protein isolation

An *M. smegmatis* strain with a 3×FLAG tag at the C terminus of subunit QcrB was generated with the oligonucleotide-mediated recombineering followed by Bxb integrase targeting (ORBIT) method (Murphy et al., 2018). This method requires transformation of the parent strain with a plasmid encoding the Che9c phage RecT annealase and Bxb1 integrase, a payload plasmid with the desired insert, and an oligonucleotide that guides integration of the payload into the chromosomal DNA. The parent strain MC2155 was transformed with plasmid pKM444, which encodes the Che9c annealase and Bxb1 integrase. The resulting strain was subsequently transformed with payload plasmid pSAB41, which encodes a 3×FLAG tag and was described previously (Guo et al., 2021), as well as the targeting oligonucleotide 5′- CAAGTCGCTCACGGCGCTCAAGGAGCACCAGGACCGCATCCACGGCAACGGGGAGACCAACGGTCATCACGGTTTGTCTGGTCAACCACCGCGGTCTCAGTGGTGTACGGTACAAACCTGATCGCTGAGATACTCGGATCGCCGCAATTCCTCTTCGGAGGGGTTGCGGCGATCTTTTTATGTGCGCTTTC-3′. The resulting strain “M. smegmatis QcrB-3xFLAG” was selected with hygromycin (50 μg/ml) and correct insertion of the 3×FLAG sequence was confirmed by colony PCR.

*M. smegmatis* was cultured in 7H9 medium (Sigma) supplemented with TDS (10 g/L tryptone, 2 g/L dextrose, 0.8 g/L NaCl). A preculture in liquid medium (15 mL) was inoculated with cells from an agar plate and grown at 30 °C with shaking at 180 rpm for 48 h. This culture was used to inoculate a larger culture (6 L), which was grown at the same conditions for a further 48 h. Cells were harvested by centrifugation at 6,900 ×g for 20 min and frozen in liquid nitrogen for subsequent use. After thawing, cells were broken by four passes through a continuous flow cell disruptor (Avestin) at 20 kpsi and membranes were harvested by centrifugation at 39,000 ×g for 30 min.

To purify CIII_2_CIV_2_, membranes were resuspended in lysis buffer (50 mM Tris-HCl pH 7.5, 100 mM NaCl, 0.5 mM EDTA) at 4 ml/g and solid dodecyl maltoside (DDM) detergent was added to 1 % (w/v) with stirring at 4 °C for 45 min. Insoluble material was removed by centrifugation at 149,000 ×g for 45 min and the solubilized protein was loaded onto a gravity column of 2 mL M2 anti-FLAG affinity matrix (Sigma). The column was washed with 10 mL of wash buffer (50 mM Tris-HCl pH 7.4, 150 mM NaCl, 0.02% [w/v] DDM) and eluted with 5 mL of wash buffer supplemented with 3×FLAG peptide at 150 μg/mL.

### Activity assays

2,3-dimethyl-[1,4]naphthoquinol (DMW) (Enamine) at 20 mM in anhydrous ethanol (400 μL) on ice was reduced with a few grains of NaBH_4_ and the reaction was quenched by addition of 4 NHCl (10 to 20 μL). Oxygen-reduction assays were performed with a Clark-type electrode (Oxygraph, Hansatech) in 1 mL of reaction buffer (50 mM Tris-HCl pH 7.5, 100 mM NaCl, 0.5 mM EDTA, and 500 nM bovine SOD [Sigma]). CIII_2_CIV_2_ was added (65 nM) and reactions were initiated by addition of 100 μM DMWH_2_. For inhibition studies, telacebec (DC Chemicals) at varying concentrations was incubated with 65 nM CIII_2_CIV_2_ in the reaction buffer for 3 h at 4 °C. This mixture was added to the Oxygraph and reactions were initiated by addition of 100 μM DMWH_2_. To account for any background oxygen reduction that still occurs in the presence of SOD, the rate of oxygen reduction in the presence of DMWH_2_ and SOD, but in the absence of CIII_2_CIV_2_, was subtracted from the rate in the presence of DMWH_2_, SOD and CIII_2_CIV_2_. The resulting oxygen reduction rates for CIII_2_CIV_2_ at different concentrations of telacebec were fit with a Python script. Individual inhibition curves, which were produced on different days with different preparations of reagent were fit individually, with the average of the IC_50_ values reported and the standard deviation of the fitted IC_50_ values reported as the error (Dahlin et al., 2004). Plots were produced using the Python matplotlib library.

### CryoEM specimen preparation and imaging

For cryoEM of inhibitor-free CIII_2_CIV_2_, enzyme at ~16 mg/mL (2 μL) was applied to homemade nanofabricated holey gold grids (Marr et al., 2014), which had previously been glow-discharged in air for 120 s at 20 mA (PELCO easiGlow), within a Vitrobot Mark III (FEI) at 4 °C and 100 % relative humidity. Grids were blotted for 24 s before freezing. For cryoEM of telacebec-bound CIII_2_CIV_2_, DMWH_2_ in ethanol was added to 100 μM (0.02% ethanol) and telacebec in DMSO was added to 25 μM (1.5% DMSO) to a solution containing purified CIII_2_CIV_2_ at ~0.08 mg/mL (6 mL). The solution was concentrated ~100-fold by centrifugation at 700 ×g with a 100 kDa molecular weight cutoff centrifuged concentrator device (Sigma). The sample (2 μL) was then applied to homemade nanofabricated holey gold grids, which had previously been glow-discharged in air for 120 s at 20 mA, within an EM GP2 (Leica) grid freezing device at 4 °C and 100 % relative humidity. Grids were blotted for 1 s before freezing.

Screening of specimens was done with an FEI Tecnai F20 electron microscope equipped with a K2 Summit direct detector device camera. High-resolution cryoEM data were collected with a Titan Krios G3 electron microscope (Thermo Fisher Scientific) operated at 300 kV and equipped with a Falcon 4 direct detector device camera. Automated data collection was done with the EPU software package. The inhibitor-free dataset consisted of 2,793 movies and telacebec-bound sample consisted of 4,308 movies. Movies were collected at a nominal magnification of 75,000× with a calibrated pixel size of 1.03 Å. Movies consisted of 30 exposure fractions over 7.7 s. The camera exposure rate and the total exposure were 5.99 e^−^/pixel/s and ~43.5 e^−^/Å^2^, respectively (**Table 1**). Contrast transfer function estimation, particle selection, motion correction, map calculation, and three-dimensional variability analysis (3DVA) were all done within the *cryoSPARC* software package (Punjani et al., 2020, 2017; Punjani and Fleet, 2021; Rubinstein and Brubaker, 2015).

### Image analysis and atomic model building

All image analysis was performed within the *cryoSPARC* software package, ver. 3 (Punjani et al., 2017), including individual particle motion correction (Rubinstein and Brubaker, 2015), non-uniform refinement (Punjani et al., 2020), and 3D Variability analysis (Punjani and Fleet, 2021). Image analysis and 3D reconstruction for each dataset was performed in the same way. Motion was corrected and contrast transfer function (CTF) parameters were estimated for each movie in patches. Manual particle selection and 2D classification was used to generate templates, which were in turn used to select of 387,777 and 1,037,709 particle images for the telacebec-bound and inhibitor-free datasets, respectively. Datasets were cleaned with 2D classification, 3D classification, and heterogeneous refinement to 70,818 and 150,885 particle images for the telacebec-bound and inhibitor-free datasets, respectively. Beam-tilt was corrected and each map was refined with non-uniform refinement without symmetry enforced. CTF values were then refined, the detergent micelle subtracted, and alignment parameters adjusted with local refinement with C2 symmetry enforced, yielding maps at 3.0 Å resolution for each dataset. 3D variability analysis was done on a pooled dataset with masks including the SOD subunit or cyt. *cc* domain. Atomic models were constructed starting from previous models of the complex (Gong et al., 2018; Wiseman et al., 2018). Additions to the models were made in *Coot* (Emsley et al., 2010) and refined with *Phenix* (Liebschner et al., 2019) and *ISOLDE* (Croll, 2018).

### Structure-Activity relation analysis studies

Insight into protein-inhibitor interaction was facilitated by analysis with the *Schrödinger* software package (Release 2019-1). The protein preparation wizard within *Schrödinger* was used to prepare the protein for modelling. Briefly, the QcrA and QcrB chains and the ligand from the PDB file were merged and pre-processed to add missing hydrogen atoms, fill in missing side chains, and adjust ionization and tautomeric states of the ligand. The hydrogen bond network between the protein’s amino acids and the ligand was optimized by allowing reorientation of amino acid side chains like His, Asn, Asp, Glu, and Gln and the ionization and tautomeric states of these side chains were estimated (Olsson et al., 2011). The resulting structure was refined to remove clashes and optimize geometry with the OPLSe* force field (Roos et al., 2019). These changes did not noticeably affect the fit of the model within the experimental cryoEM density map.

## Data availability

Data deposition: all electron cryomicroscopy maps described in this article have been deposited in the Electron Microscopy Data Bank (EMDB) (accession numbers EMD-XXXX to EMD-XXXX) and atomic models have been deposited in the Protein Database (PDB) (accession numbers XXXX).

## Author contributions

PB and JLR conceived the project and coordinated experiments. PB, PI, and JLR supervised the research. SAB prepared the *M. smegmatis* strain used for these experiments. DJY cultured cells, purified protein, prepared and imaged cryoEM specimens. DJY and SK performed enzyme assays. DJY and JDT performed cryoEM image analysis and atomic model building and refinement. JDT and RA analyzed the telacebec structure-activity relations. DJY, JDT, and JLR wrote the manuscript and prepared the figures with input from the other authors.

## Acknowledgements

DJY was supported by a Canada Graduate Scholarship from the Canadian Institutes of Health Research (CIHR) and a Queen Elizabeth II Graduate Scholarship in Science and Technology from the University of Toronto Department of Medical Biophysics, JMDT was supported by a postdoctoral fellowship from the Canadian Institutes of Health Research, and JLR was supported by the Canada Research Chairs program. This research was supported by CIHR grant JT162186 (JLR), The Alice and Knut Wallenberg Foundation grant 2019.0043 (PB), and Swedish Research Council grant 2018-04619 (PB). Cryo-EM data were collected at the Toronto High-Resolution High-Throughput Cryo-EM facility, supported by the Canada Foundation for Innovation and Ontario Research Fund.

## Figure Supplements

**Figure 1 – Figure Supplement 1.**
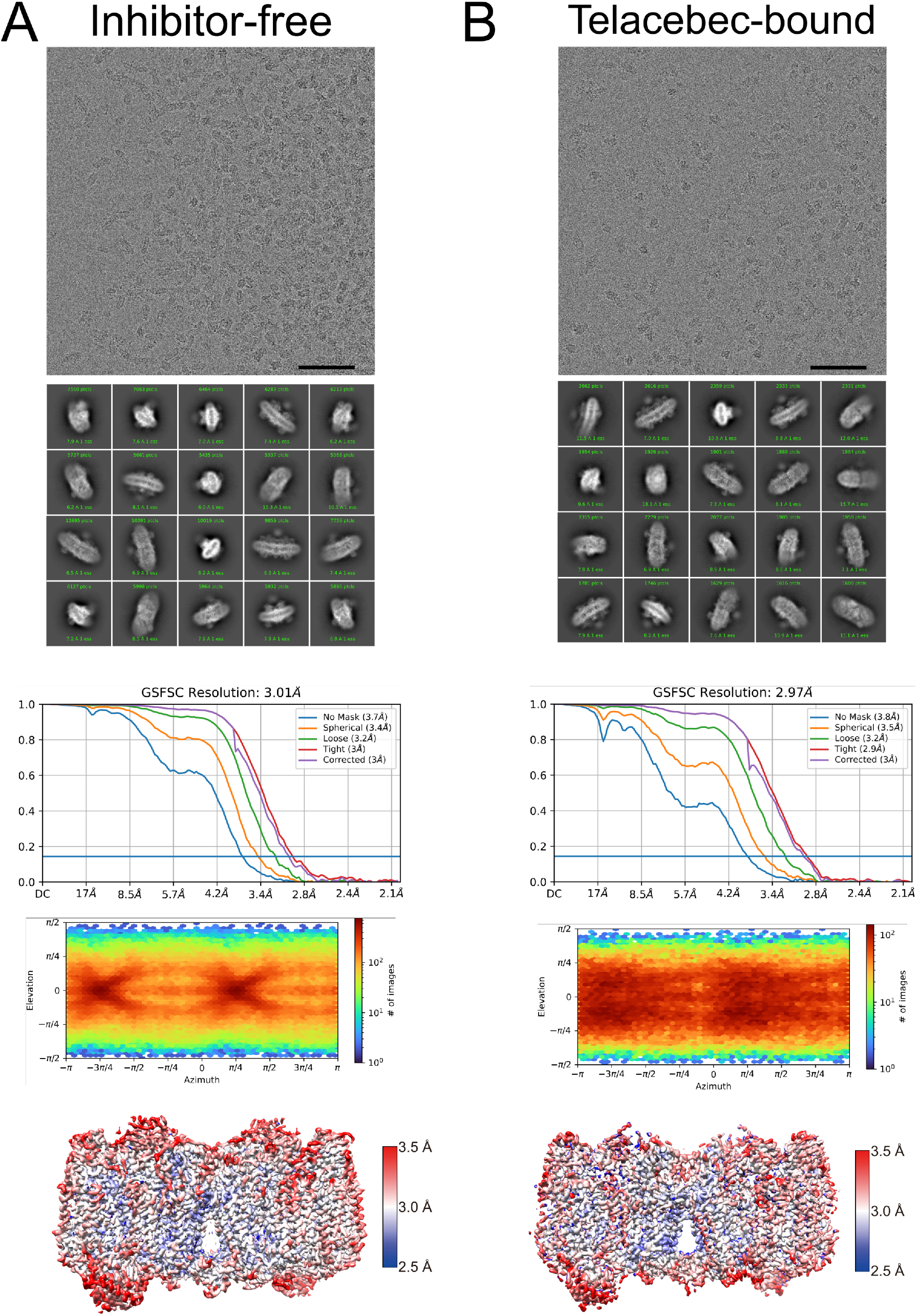
CryoEM map validation. **A,** An example micrograph, class average images, Fourier Shell Correlation curves (including correction for the effects of masking following gold-standard refinement), orientation distribution plot, and local resolution estimate are shown for the inhibitor-free CIII_2_CIV_2_ map. **B**, An example micrograph, class average images, Fourier Shell Correlation curves (including correction for the effects of masking following gold-standard refinement), orientation distribution plot, and local resolution estimate are shown for the telacebec-bound CIII_2_CIV_2_ map. Scale bars, 50 nm.

**Figure 1 – Figure Supplement 2.**
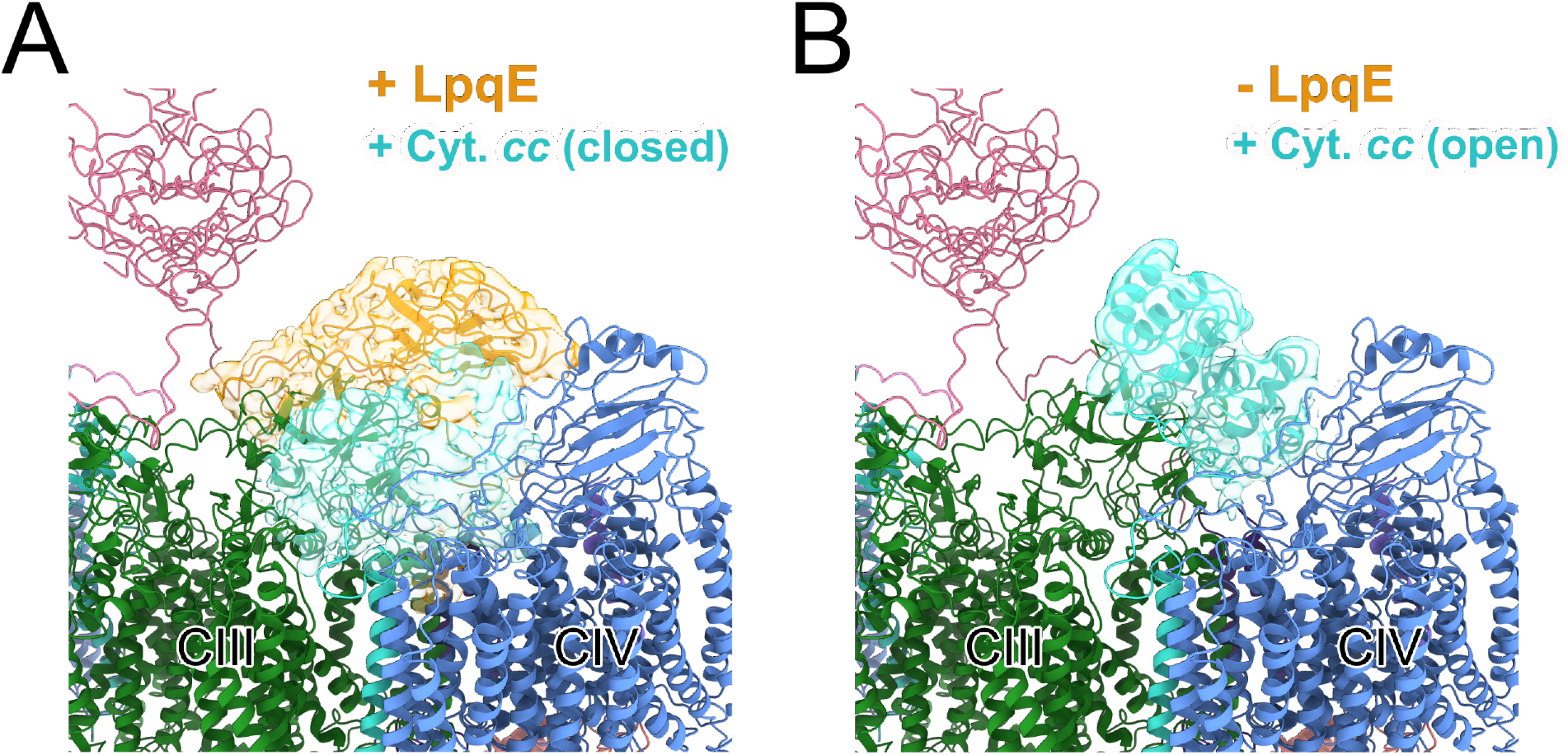
3D variability analysis of LpqE and the cyt. *cc* domain. Focusing on one half of the CIII_2_CIV_2_ supercomplex with symmetry expansion to allow all half complexes to contribute, complexes were identified with LpqE (*gold*) and the cyt. *cc* domain (*cyan*) in the ‘closed’ position (**A**) and without LpqE and the cyt. *cc* domain (*cyan*) in the ‘open’ position (**B**).

**Figure 1 – Figure Supplement 3.**
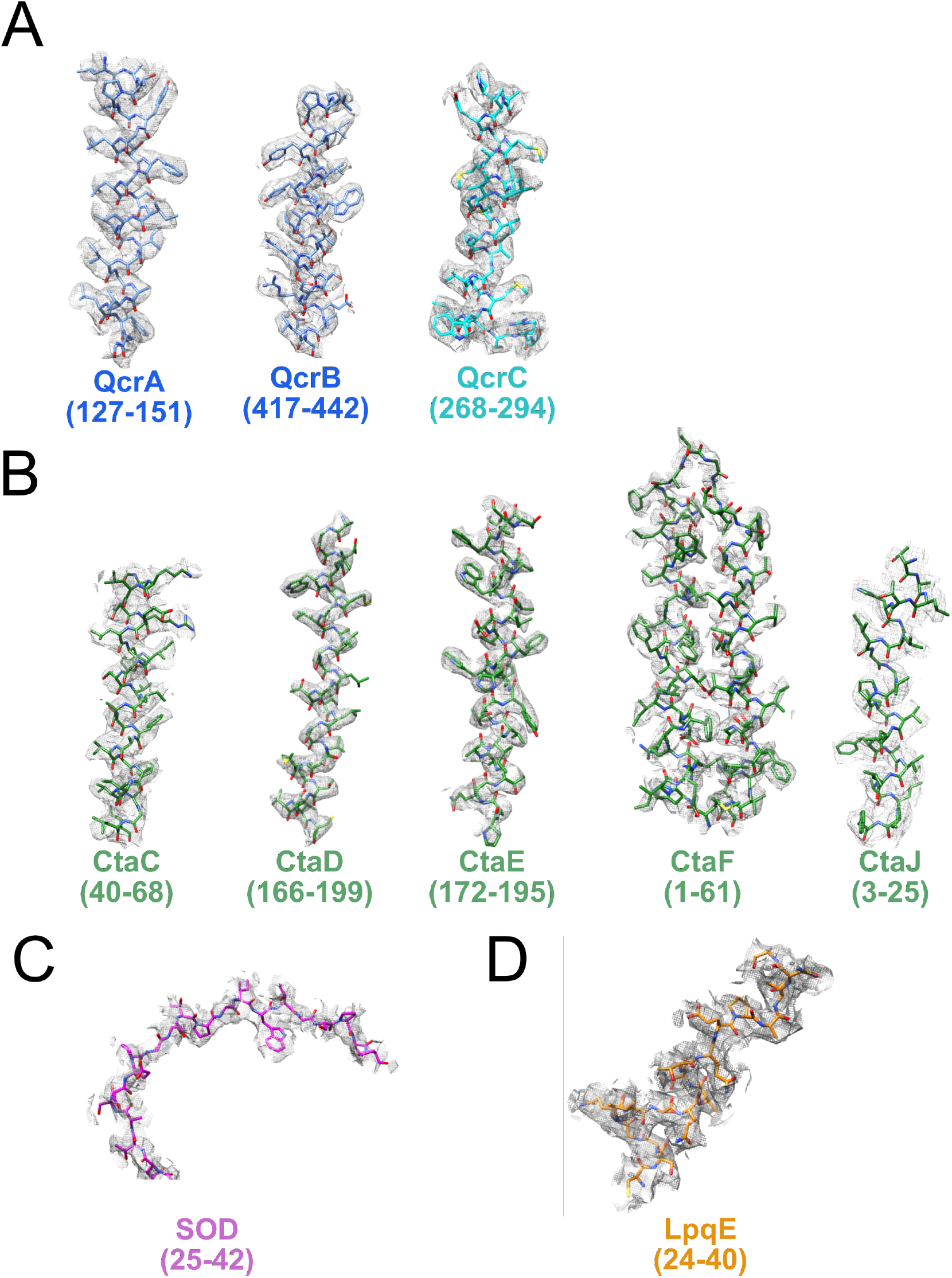
Examples of model in map fit. Examples for model in map fit are shown for the CIII part of the supercomplex (**A**), the CIV part of the supercomplex (**B**), as well as the SOD (**C**) and LpqE (**D**) subunits.

**Figure 3 – Figure Supplement 1.**
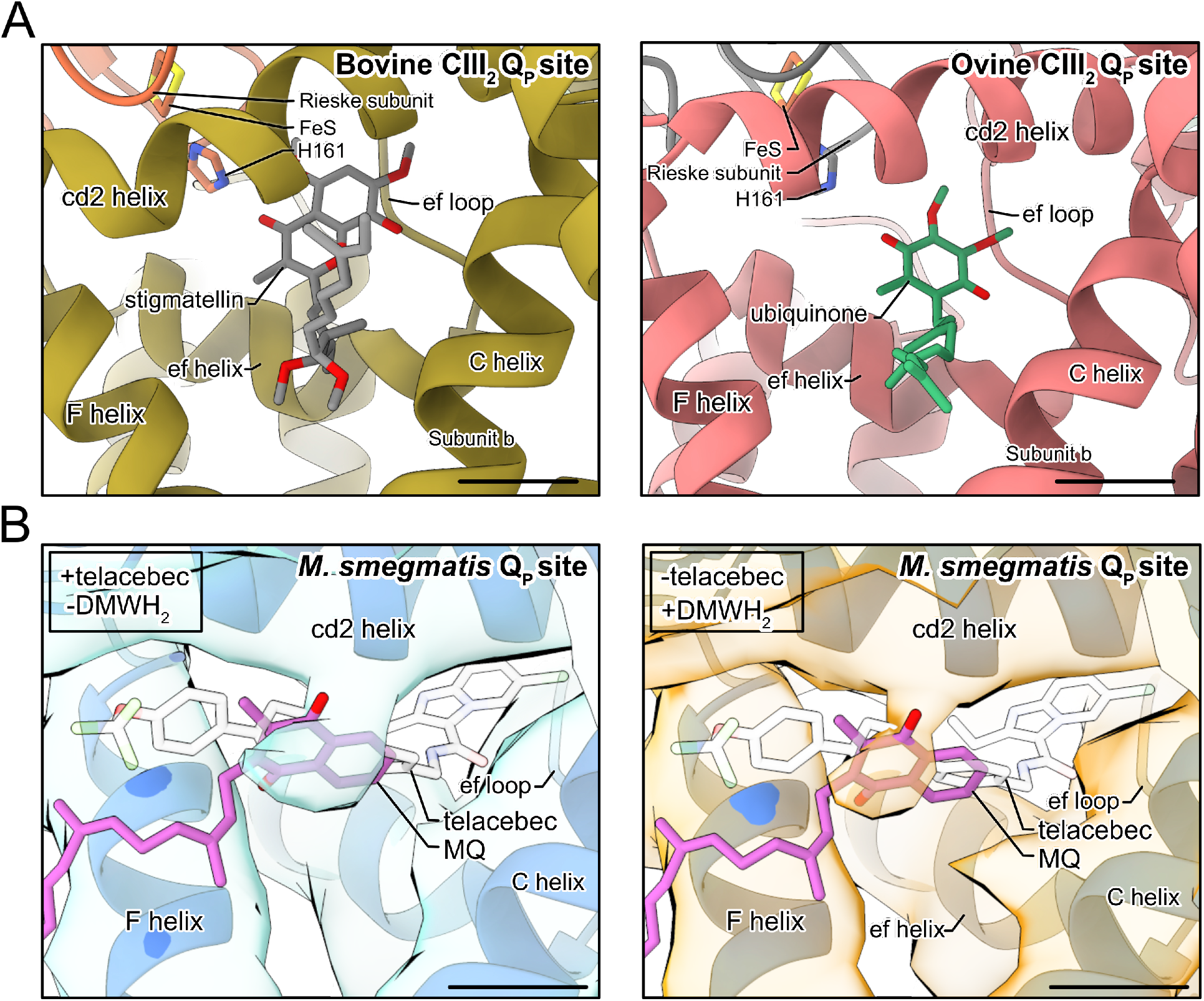
Binding of small molecules in the Q_P_ site of CIII. **A**, Both stigmatellin (*grey model*) in the bovine CIII_2_ Q_P_ site (*left*) (PDB: 1PPJ) (Huang et al., 2005) and ubiquinone (*green model*) in the ovine CICIII_2_ supercomplex Q_P_ (*right*) (PDB: 6Q9E) (Letts et al., 2019) bind deep in the site close to the FeS group, different from endogenous MQ with *M. smegmatis* CIII_2_CIV_2_. **B,** CryoEM density from the 200 kV screening microscope shows that CIII_2_CIV_2_ with telacebec but without DMWH_2_ at 4.7 Å resolution (*left*) and with DMWH_2_ but without telacebec at 4.4 Å resolution (*right*) is consistent with binding of MQ (*pink model*) but not telacebec (*white model*) in the Q_P_ site. Scale bars, 5 Å.

